# Microbial persistence beneath temporal restructuring in university wastewater metagenomes

**DOI:** 10.64898/2026.10.13.758942

**Authors:** Gabriela Franco, Archana Anand

## Abstract

Temporal changes in wastewater microbiomes can reflect shifts in microbial abundance, detectable membership, and genome recovery. We paired genome-resolved and read-based metagenomics to distinguish these signals in 25 spatially pooled university wastewater samples collected during wet and dry periods. The workflow recovered 203 wet-period and 230 dry-period metagenome-assembled genomes (MAGs). Community composition differed between periods in both MAG-derived presence/absence profiles (Jaccard PERMANOVA, R^2^ = 0.548, p = 0.001) and read-based relative-abundance profiles (Bray–Curtis PERMANOVA, R^2^ = 0.167, p = 0.001). Greater dry-period read-based richness persisted after rarefaction but disappeared under more stringent abundance thresholds, indicating a substantial contribution from low-abundance detections. Of 57 period-associated core species in the MAG-derived profiles, 14 had exact species-level matches in the read-based dataset. All 14 remained detectable at ≥90% prevalence in the opposite period, and 13 had higher relative abundance in the period corresponding to their MAG-based association. Continued species detection therefore accompanied changes in community composition and MAG-based representation. Perfect/Strict CARD/RGI resistance-associated annotations occurred in 94.1% of wet-period and 92.2% of dry-period MAGs and were distributed across shared-core and period-associated taxa. These findings reveal repeated detection of a matched subset of microbial taxa beneath temporal community restructuring, together with widespread resistance-associated genetic potential among recovered genomes. Combining read-based and genome-resolved profiles helps distinguish changes in detectable membership from changes in the genomic characterization available for wastewater surveillance.

**Importance:** A changing wastewater microbial profile can suggest that organisms have disappeared or been replaced. In this university wastewater time series, community composition differed between sampling periods, yet all 14 species that could be matched between the period-associated MAG core and read-based profiles remained highly prevalent in the opposite period. Most also shifted in relative abundance in the direction of their MAG-based association. Thus, continued species detection can coexist with changes in community composition and genome representation. Resistance-associated annotations were widespread among the recovered genomes, providing complementary information about their genetic potential. Paired analyses can help wastewater surveillance distinguish organisms that remain detectable from those for which genomic context is available.

## INTRODUCTION

Wastewater metagenomics characterizes microbial communities and antimicrobial resistance (AMR) potential across human, environmental, and sewer-associated inputs (Pruden et al., 2021; Fierer et al., 2022). Interpreting changes in these profiles requires distinguishing shifts in relative abundance from changes in detectable membership. A further distinction arises in genome-resolved analyses: changes in the genomes that can be reconstructed or detected may alter the apparent representation of taxa without demonstrating their disappearance. Separating these signals is important for interpreting microbial dynamics and the genomic information available for surveillance.

Read-based and genome-resolved analyses offer complementary views of a wastewater community. Read-based taxonomic profiling estimates composition from sequence assignments, whereas genome-resolved metagenomics reconstructs genomes that can place resistance-associated annotations in microbial context. Assembly and binning require sufficient sequence information, so read-level evidence for a species may be available even when a corresponding metagenome-assembled genome (MAG) is not recovered. Sequencing depth also affects inferred taxonomic and AMR profiles (Gweon et al., 2019). A taxon associated with one period in MAG-derived profiles may therefore remain detectable by read-based profiling in another period.

Previous campus wastewater research has documented spatial and temporal variation in microbial communities and resistance-gene profiles, including substantial contributions from environmental organisms (Fierer et al., 2022). Longer sewage time series have distinguished seasonal, human-derived, and environmental microbial components, with implications for source-attributed surveillance (Becsei et al., 2024). These studies establish the importance of community dynamics. A complementary question is whether taxa associated with one temporal window in genome-resolved profiles remain detectable when the same samples are examined through read-based profiling.

Repeated sampling of university wastewater provides a tractable setting for this comparison. Passive tampon samplers integrate material over a deployment interval and have been used for wastewater surveillance (Bivins et al., 2022; West et al., 2023). Here, wet and dry sampling windows provided contrasting periods within one campus sewer network. Because these windows also differed in calendar time and potentially campus occupancy, we examine period-associated variation rather than attributing differences specifically to rainfall or season.

We paired genome-resolved and read-based analyses of San Francisco State University wastewater to examine community restructuring and continued taxon detection across wet and dry sampling periods. We compared community composition and richness, then tested whether period-associated core species in MAG-derived profiles remained detectable in the read-based dataset. We also examined pathogen-associated taxonomic signals and the distribution of resistance-associated annotations among recovered genomes. Together, these analyses connect temporal community patterns with the taxonomic and genomic information available for wastewater surveillance.

## METHODS

### Wastewater sampling and sample processing

Wastewater was sampled at three locations within the San Francisco State University campus sewer network during December 2024–February 2025 (wet period) and April–August 2025 (dry period). Samples were collected approximately weekly at a consistent time of day using a passive, Moore-swab-inspired approach. Commercially available Tampax Super tampons were suspended directly in the wastewater stream for 24 h, consistent with previous applications of passive wastewater sampling (Bivins et al., 2022; West et al., 2023). The study comprised 25 spatially pooled samples: 11 from the wet period and 14 from the dry period.

Following collection, wastewater recovered from the three sites was pooled in equal volumes. Where sufficient material was available, 12.5 mL from each site was combined; for lower-volume collections, 10 mL from each site was used. The resulting composite was divided into aliquots for downstream processing. Nucleic acids were concentrated and extracted using the Nanotrap Microbiome A and B Enviro Water Protocol with the NucleoMag kit (APP-081) or the Nanotrap Microbiome Combined 10-mL Automated Protocol with the NucleoMag kit on a KingFisher Apex platform (APP-076).

### Shotgun metagenomic sequencing

Extracted DNA was submitted to Zymo Research for shotgun metagenomic sequencing at a target depth of approximately 80 million paired-end reads per sample. Libraries were prepared using the Illumina DNA Library Prep Kit with up to 500 ng DNA, unique dual-index 10-bp barcodes, and Nextera adapters, following the manufacturer’s protocol. The ZymoBIOMICS Microbial Community DNA Standard was included as a positive control for each library preparation. Sequencing was performed on an Illumina NovaSeq platform.

### Genome-resolved metagenomic analysis

Raw reads were quality filtered using BBDuk from BBTools (v39.33) with adapter and PhiX removal (ref=adapters,phix), right-end k-mer trimming (ktrim=r, k=23, mink=11, hdist=1), paired-read overlap trimming (tpe, tbo), and quality trimming of both read ends at a Phred score threshold of 10 (qtrim=rl, trimq=10); reads shorter than 50 bp after trimming were discarded (minlen=50). Filtered reads were assembled using MEGAHIT (v1.2.9) (Li et al., 2015). Metagenome-assembled genomes (MAGs) were recovered using MaxBin2 (v2.2.7) (Wu et al., 2016) and evaluated using CheckM (v1.2.4) (Parks et al., 2015). MAGs with estimated completeness ≥50% and contamination ≤10% were retained. These thresholds yielded 203 MAGs from wet-period samples and 230 from dry-period samples. Taxonomy was assigned using GTDB-Tk (v2.6.1; GTDB release R226; Chaumeil et al., 2022). Reads were mapped to recovered MAGs using Bowtie2 (v2.5.5; Langmead and Salzberg, 2012) with the --very-sensitive preset. Mean coverage and covered fraction were calculated using CoverM (v0.7.0; Aroney et al., 2025), and a MAG was considered detected in a sample when it had ≥1× mean coverage and ≥30% genome coverage breadth.

Species assigned by GTDB-Tk were classified using a ≥90% within-period prevalence threshold in the MAG-derived profiles. Shared-core species met this threshold in both periods. Wet-core-only and dry-core-only species met the threshold in the indicated period but not in the other; together, these are termed period-associated core species. In this manuscript, reconstruction refers to assembly and binning, whereas MAG detection refers to read mapping that meets the coverage thresholds above. Core membership describes prevalence in the MAG-derived profiles.

### Read-based taxonomic profiling

Quality-controlled reads were classified using Kraken2 (v2.17.1; Wood et al., 2019) with the Standard-16 database dated 2026-06-26. Species-level abundance estimates were generated using Bracken (v3.1; Lu et al., 2017). Human-classified reads were removed before microbial community analyses, and relative abundances were renormalized. These profiles were used to estimate species richness and abundance-weighted community composition. Period-associated core species in the MAG-derived profiles were compared with the Bracken dataset using exact species-name matches. Within the matched subset, read-based prevalence and relative abundance were compared between periods; unmatched species remained unresolved.

### Pathogen-associated organisms

Pathogen-associated taxa were identified from documented associations with human disease in WHO, CDC, NCBI, and FDA resources and the clinical literature (WHO, 2024; NCBI, n.d.-a, n.d.-b; CDC, 2024, 2025, 2026; FDA, 2012; Elsaghir and Reddivari, 2023; Kamali et al., 2024). Reporting was restricted to taxa with mean relative abundance ≥0.01% and prevalence ≥50% in at least one period. ESKAPE screening considered five species—*Enterococcus faecium, Staphylococcus aureus, Klebsiella pneumoniae, Acinetobacter baumannii*, and *Pseudomonas aeruginosa*—and the genus-level group *Enterobacter* spp. These assignments identify pathogen-associated taxa rather than pathogenic strains or viable organisms.

### AMR annotation

Recovered MAGs were analyzed using the Resistance Gene Identifier (RGI v6.0.5) against the Comprehensive Antibiotic Resistance Database (CARD v3.2.7; Alcock et al., 2023). The initial output included Perfect, Strict, and Loose predictions, of which 99.6% were Loose. Principal AMR summaries were restricted to Perfect and Strict calls and described the frequency, annotation counts, and database-assigned resistance categories among recovered MAGs. These annotations represent resistance-associated genetic potential.

### Statistical analysis

Statistical analyses were performed in R, with community analyses conducted using vegan. Two-sided Wilcoxon rank-sum tests compared per-sample MAG counts, MAG-derived species richness, and read-based species richness between periods. Wilcoxon tests also compared the relative abundances of matched period-associated core species and annotation counts per MAG. Sampling dates were the units of comparison for community analyses; per-MAG annotation comparisons described the recovered genome collections. Read-based richness was evaluated after rarefaction to a common Bracken count depth and across minimum relative-abundance thresholds.

Community composition was compared using PERMANOVA with 999 permutations. MAG-derived species presence/absence profiles were analyzed using Jaccard dissimilarities, and Bracken relative-abundance profiles were analyzed using Bray–Curtis dissimilarities after human-read removal and renormalization. Principal coordinates analysis (PCoA) visualized community dissimilarities. Multivariate dispersion was evaluated using betadisper and permutation tests with 999 permutations. Sensitivity analyses examined exclusion of low-coverage or low-classification samples and restriction of MAG-derived richness comparisons to species meeting prevalence thresholds. Statistical significance was evaluated at p < 0.05.

## RESULTS

### Wet- and dry-period samples differed in MAG representation and richness

The genome-resolved workflow yielded 203 wet-period and 230 dry-period MAGs meeting the completeness and contamination thresholds. Per-sample MAG counts were greater in the dry period (median 218 versus 177; Wilcoxon W = 143, p = 3.28 × 10□□). This difference remained after exclusion of low-coverage samples (W = 130, p = 6.10 × 10□□).

Bacillota dominated both MAG collections, accounting for 70.4% of wet-period and 63.5% of dry-period MAGs, followed by Pseudomonadota and Bacteroidota (Figure 1). Differences were also evident among less represented phyla, including Patescibacteriota and Campylobacterota in the dry-period collection.

**Figure 1.**
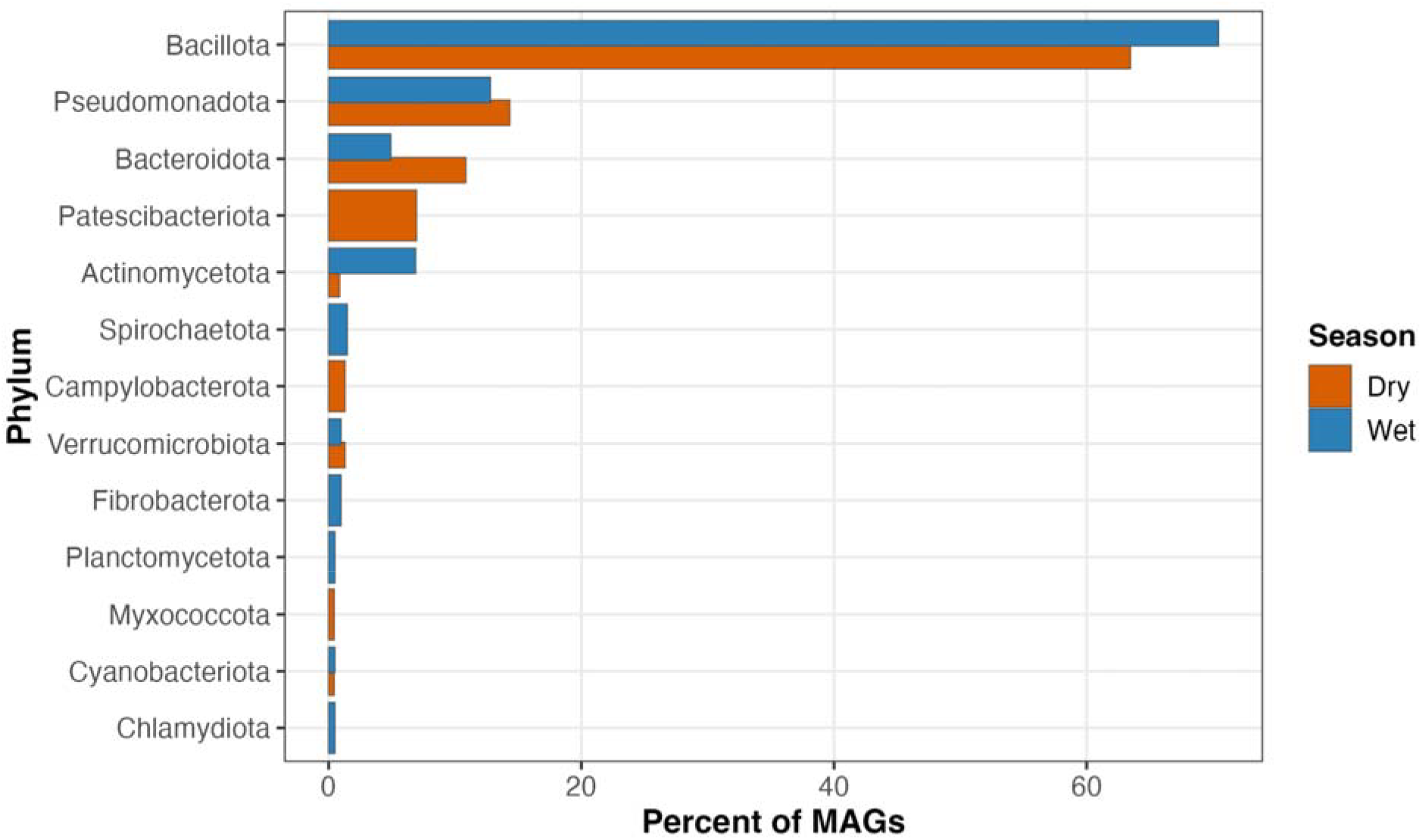
Phylum composition of the wet- and dry-period MAG collections. Bars show the percentage of recovered MAGs assigned to each phylum in the wet (n = 203 MAGs) and dry (n = 230 MAGs) collections. Percentages describe genome counts within each collection and are not weighted by sample-level relative abundance.

### Period-associated differences extended to MAG-derived species composition

MAG-derived species richness was greater in dry-than wet-period samples (median 83 versus 62 species; W = 143, p = 3.20 × 10□ □). The difference remained after removal of low-coverage samples and when the analysis was restricted to species present in ≥50% of samples, but was not significant at a ≥75% prevalence threshold. The richness contrast therefore depended on the prevalence threshold used to define the species set.

MAG-derived species composition differed between sampling periods based on Jaccard distances (PERMANOVA, F = 27.8, R^2^ = 0.548, p = 0.001; Figure 2). There was no evidence of unequal multivariate dispersion between periods (p = 0.645).

**Figure 2.**
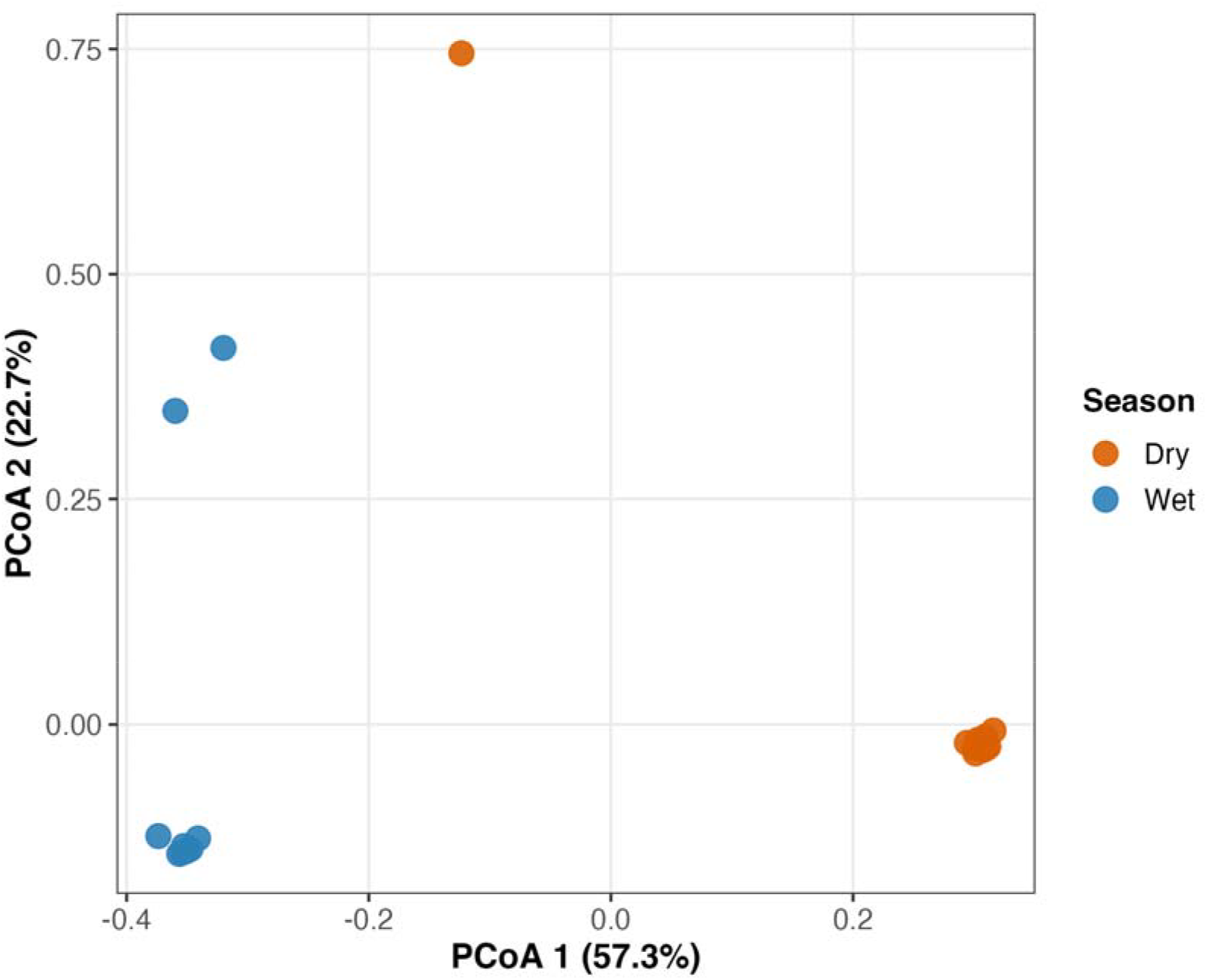
Period-associated structure in MAG-derived species profiles. PCoA of Jaccard dissimilarities calculated from species presence/absence. Points represent samples and colors indicate sampling period. Axis labels show the percentage of variation represented by each coordinate. PERMANOVA: F = 27.8, R^2^ = 0.548, p = 0.001; multivariate dispersion test: p = 0.645.

### Shared and period-associated core species in the MAG-derived profiles

At the ≥90% within-period prevalence threshold, 29 species formed a shared core, 25 were wet-core only, and 32 were dry-core only (Figure S1). Of these 57 period-associated core species, 52 were not reconstructed from the opposite period. We therefore examined whether period-associated core membership coincided with loss of read-based species detection.

### Read-based profiles distinguish community restructuring from continued species detection

Read-based relative-abundance profiles also differed between periods (Bray–Curtis PERMANOVA, F = 4.60, R^2^ = 0.167, p = 0.001; Figure 3), without evidence of unequal multivariate dispersion (p = 0.798). Observed Bracken species richness was greater in dry-period samples (median 4,770 versus 3,059; p = 0.017). This difference persisted after rarefaction but disappeared under more stringent abundance thresholds, indicating a substantial contribution from low-abundance detections. Period-level mean abundances of dominant read-based taxa are shown in Figure S2.

**Figure 3.**
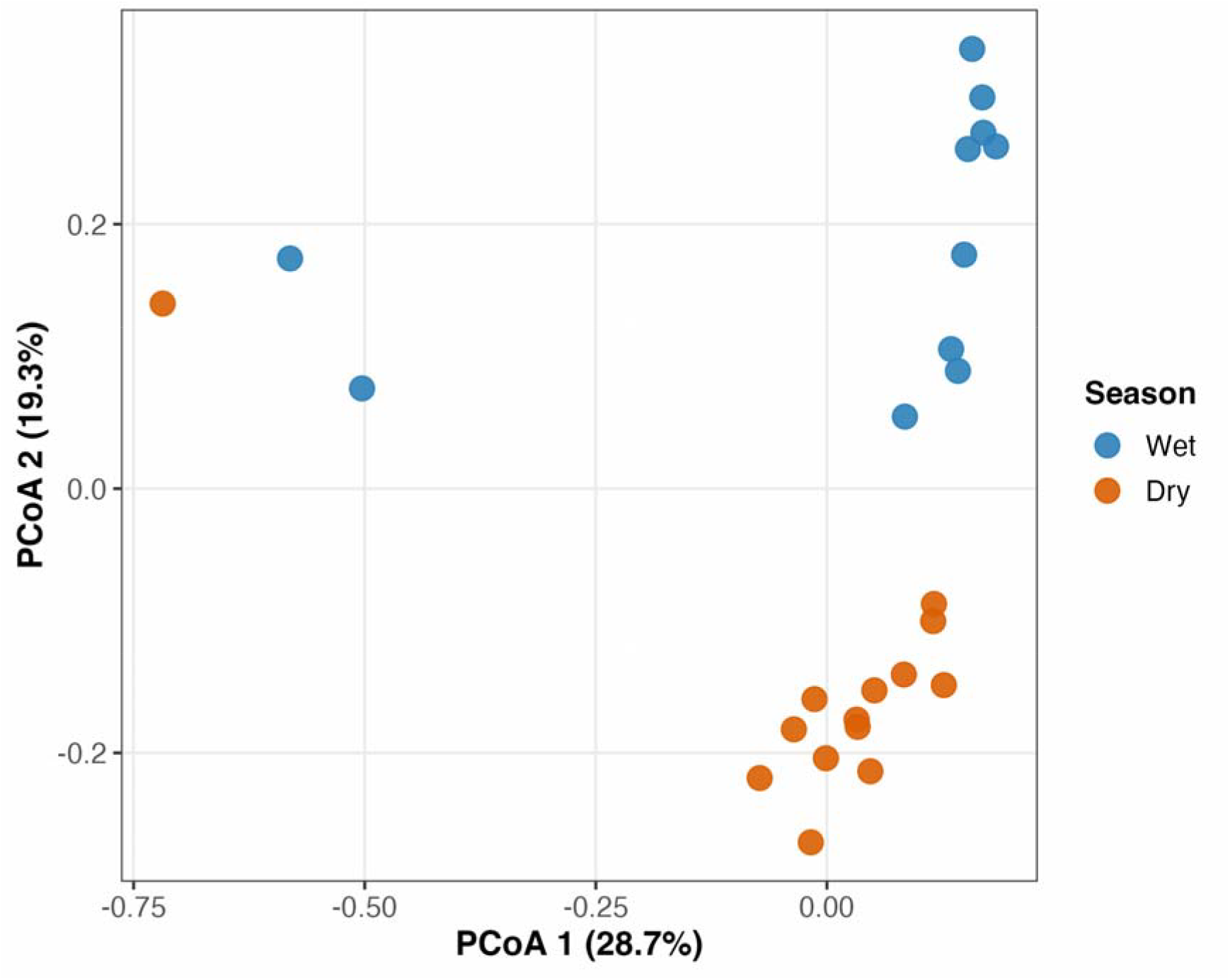
Period-associated structure in read-based species profiles. PCoA of Bray–Curtis dissimilarities calculated from Bracken relative abundances after human-read removal and renormalization. Points represent samples and colors indicate sampling period. Axis labels show the percentage of variation represented by each coordinate. PERMANOVA: F = 4.60, R^2^ = 0.167, p = 0.001; multivariate dispersion test: p = 0.798.

Fourteen of the 57 period-associated core species had exact species-name matches in the Bracken dataset: seven wet-core and seven dry-core species (Table 1). All 14 were detected in ≥90% of samples from the opposite period. Six of seven wet-core species had higher read-based relative abundance in the wet period, and all seven dry-core species had higher relative abundance in the dry period. Thus, 13 of 14 matched species showed abundance differences in the direction of their MAG-based period association while remaining highly prevalent in the opposite period. The other 43 species were unresolved by exact-name matching. Continued read-based detection in the matched subset demonstrates that period-associated MAG representation did not imply species-level absence.

**Table 1.** Continued read-based detection of period-associated core species matched between taxonomic frameworks.

| MAG-based core category | Exactly matched species, n | Detected at $\geq 90\%$ prevalence in opposite period, n | Higher relative abundance in associated period, n |
| --- | --- | --- | --- |
| Wet-core only | 7 | 7 | 6 |
| Dry-core only | 7 | 7 | 7 |
| Total | 14 | 14 | 13 |

Exact species-name matches were available for 14 of 57 period-associated core species; the other 43 remained unresolved. The abundance column reports direction of the period contrast, not statistical significance.

### Pathogen-associated taxa were repeatedly detected across sampling periods

Read-based profiling identified 15 pathogen-associated species meeting the reporting thresholds (Figure 4). *Pseudomonas aeruginosa, Escherichia coli, Acinetobacter baumannii, Klebsiella pneumoniae, Salmonella enterica*, and *Aeromonas hydrophila* were detected in every sample from both periods. Their mean relative abundances nevertheless differed: *P. aeruginosa* averaged 1.89% in the wet period and 1.25% in the dry period, whereas *E. coli* averaged 0.50% and 1.62%, respectively.

**Figure 4.**
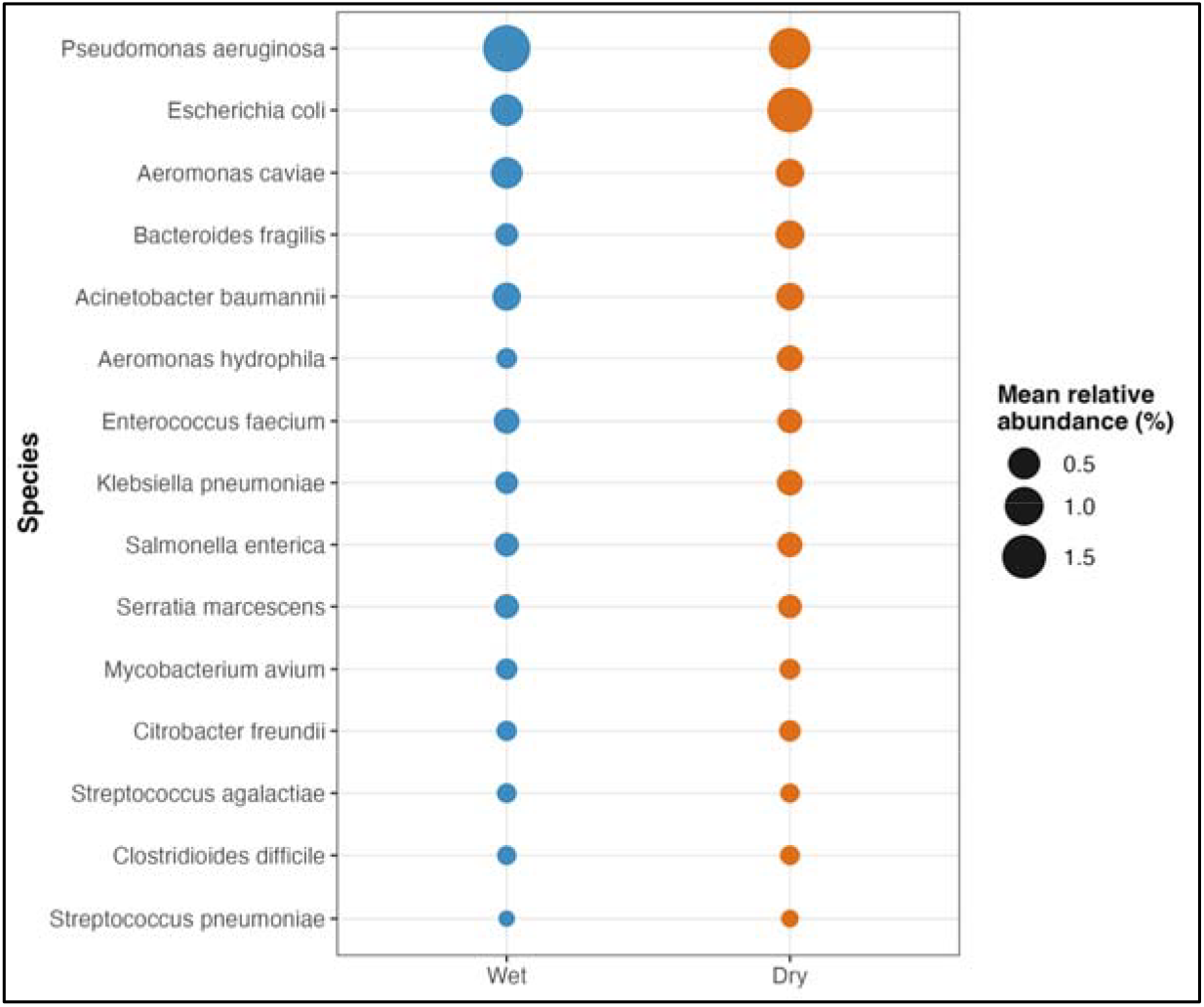
Read-based relative abundance of pathogen-associated species. Bubble size represents mean relative abundance (%) within each period. Species met mean relative abundance ≥0.01% and prevalence ≥50% in at least one period. Taxonomic detection alone does not establish viability or pathogenic strain identity.

Read-based profiles detected all six ESKAPE groups. No exact-species MAG counterparts were recovered for the five species-defined targets, although other *Acinetobacter* and *Pseudomonas* species were reconstructed. The genus-level *Enterobacter* group was excluded from this exact-species comparison. These observations indicate a difference between taxonomic detection and the availability of corresponding reconstructed genomes.

### Resistance-associated annotations were widespread among recovered genomes

Perfect or Strict CARD/RGI annotations occurred in 191 of 203 wet-period MAGs (94.1%) and 212 of 230 dry-period MAGs (92.2%). Median counts were eight and six annotations per MAG, respectively (Wilcoxon W = 22,132, p = 0.350). These summaries characterize the recovered genome collections.

Antibiotic target alteration was the dominant database-assigned mechanism among Perfect/Strict annotations in both periods. Glycopeptide-associated annotations, predominantly CARD van-family models, and tetracycline-associated annotations were prominent.

Perfect/Strict annotations occurred in shared-core taxa, including *E. coli*, and in wet- and dry-core taxa (Figure 5). Resistance-associated genetic potential was therefore distributed across shared and period-associated components of the recovered community.

**Figure 5.**
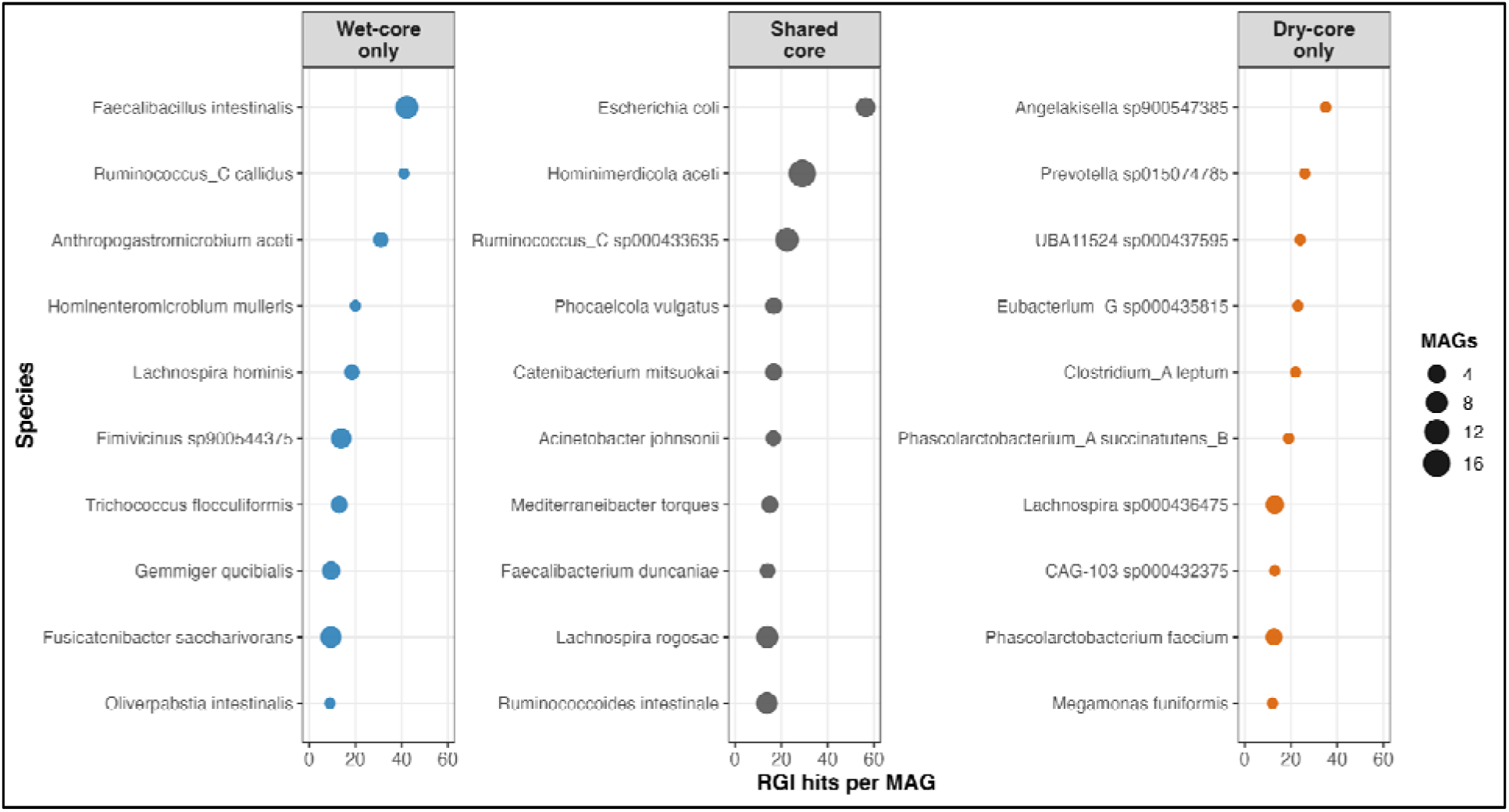
Resistance-associated annotations among selected core taxa. Horizontal position indicates RGI hits per MAG, and bubble size indicates the number of MAGs represented. Panels group taxa as wet-core only, shared core, or dry-core only, using the ≥90% within-period prevalence definition. Counts describe genomic annotations and are not weighted by sample-level host abundance.

## DISCUSSION

Temporal community restructuring coexisted with continued species detection in campus wastewater. Both MAG-derived presence/absence and read-based relative-abundance profiles distinguished the sampling periods. Within the 14 period-associated core species matched across frameworks, all remained highly prevalent in the opposite period and 13 showed abundance differences consistent with their MAG-based association. These findings extend the description of campus wastewater variability (Fierer et al., 2022) by showing how complementary profiles distinguish a change in community composition from loss of detection for a defined subset of species.

The matched-species result is consistent with abundance contributing to differences in genome representation. However, the comparison does not identify a causal recovery threshold or quantify the proportion of community-wide turnover attributable to recovery limits. Exact-name matching resolved 14 of 57 period-associated core species; the remaining 43 cannot be assigned the same behavior. Taxonomic correspondence, reference-catalog construction, sequencing depth, assembly, and binning can all influence the comparison. Persistence here means repeated species-level detection, which may reflect continued inputs as well as residence in the sewer and does not establish continuity of the same strains.

The richness analyses show that low-abundance detections contributed substantially to the greater dry-period read-based richness: the difference survived rarefaction but disappeared under stricter abundance thresholds. At the same time, abundance-weighted community composition differed between periods.

These observations support a view of temporal restructuring that includes shifts in relative abundance alongside changes in the less abundant detectable community. The two PERMANOVA R^2^ values describe different community representations and distance metrics and cannot quantify how much turnover is attributable to genome recovery. Their agreement on period-associated structure provides complementary evidence of community change.

For surveillance, the distinction between taxonomic detection and genomic characterization determines the context available for interpreting an organism’s signal. Pathogen-associated species were repeatedly detected by read-based profiling, while corresponding genomes were not always reconstructed. Genome recovery therefore limited the resistance context available for some taxonomic detections. Species assignments alone do not establish viability, pathogenic strain identity, or infection in the contributing population. Likewise, detecting a pathogen-associated species and a resistance annotation elsewhere in a metagenome does not demonstrate that the detected species carries that determinant.

Resistance-associated annotations were widespread in the reconstructed community, with Perfect/Strict calls in more than 90% of MAGs in each period and representation across shared-core and period-associated taxa. This provides genomic context for the recovered fraction of the community. Similar annotation counts per MAG, however, can coexist with differences in gene identity, host composition, and sample-level abundance. Genome completeness, size, redundancy, and taxonomic composition also affect these counts, and MAGs are not independent environmental replicates. The per-MAG comparison therefore does not establish temporal resistome stability. Individual van-family annotations additionally require model and genomic-context evaluation before inferring a complete glycopeptide-resistance system or phenotype.

The sampling design limits causal and spatial inference. Pooling material from three locations produced one composite per sampling date and prevented estimation of site-level variation. Passive sampling integrated material over 24 h without measuring the wastewater volume represented, so these profiles cannot provide absolute concentrations. The wet and dry windows also differed in calendar time and potentially campus occupancy, precluding attribution to rainfall alone. Extraction protocol, sequencing batch, and temporal dependence are additional considerations when interpreting period-associated differences. The findings therefore describe this campus time series and the matched taxonomic subset, with broader generalization requiring further validation.

## Conclusion

Campus wastewater communities differed between wet and dry periods while all 14 matched period-associated core species remained highly prevalent by read-based profiling. Abundance differences in 13 of these species accompanied their MAG-based period associations, demonstrating that community restructuring can occur alongside continued species detection. Resistance-associated annotations were widespread among recovered genomes from both periods. Integrating read-based and genome-resolved analyses provides a clearer account of which taxa remain detectable and which can be characterized genomically during longitudinal wastewater surveillance.

## Data and code availability

Sequence data are being prepared for deposition in the NCBI Sequence Read Archive, and metagenome-assembled genomes will be deposited in GenBank. Supporting sample metadata, processed data tables, and analysis scripts will be deposited in a public repository. Accession numbers and repository links will be provided in the revised manuscript, and all deposited materials will be publicly accessible no later than publication. Data and code supporting the reported findings will be made available to editors and reviewers upon request during peer review.

## Acknowledgments and funding

We thank the College of Science and Engineering, Department of Biology, and Facilities at San Francisco State University for their support. This work was supported by the National Science Foundation Center for Cellular Construction (DBI-1548297) and the U.S. Department of Energy, Office of Science, Office of Biological and Environmental Research, through Phage Pathways (DE-SC0025673) and the Biopreparedness Research Virtual Environment (BRaVE) Phage Foundry at Lawrence Berkeley National Laboratory (DE-AC02-05CH11231).

## Author contributions

Archana Anand acquired funding, provided resources, designed the study, and contributed to manuscript writing. Gabriela Franco conducted field sampling and laboratory work, performed data analysis, and contributed to manuscript writing.

## Supplementary Figures

**Figure S1.**
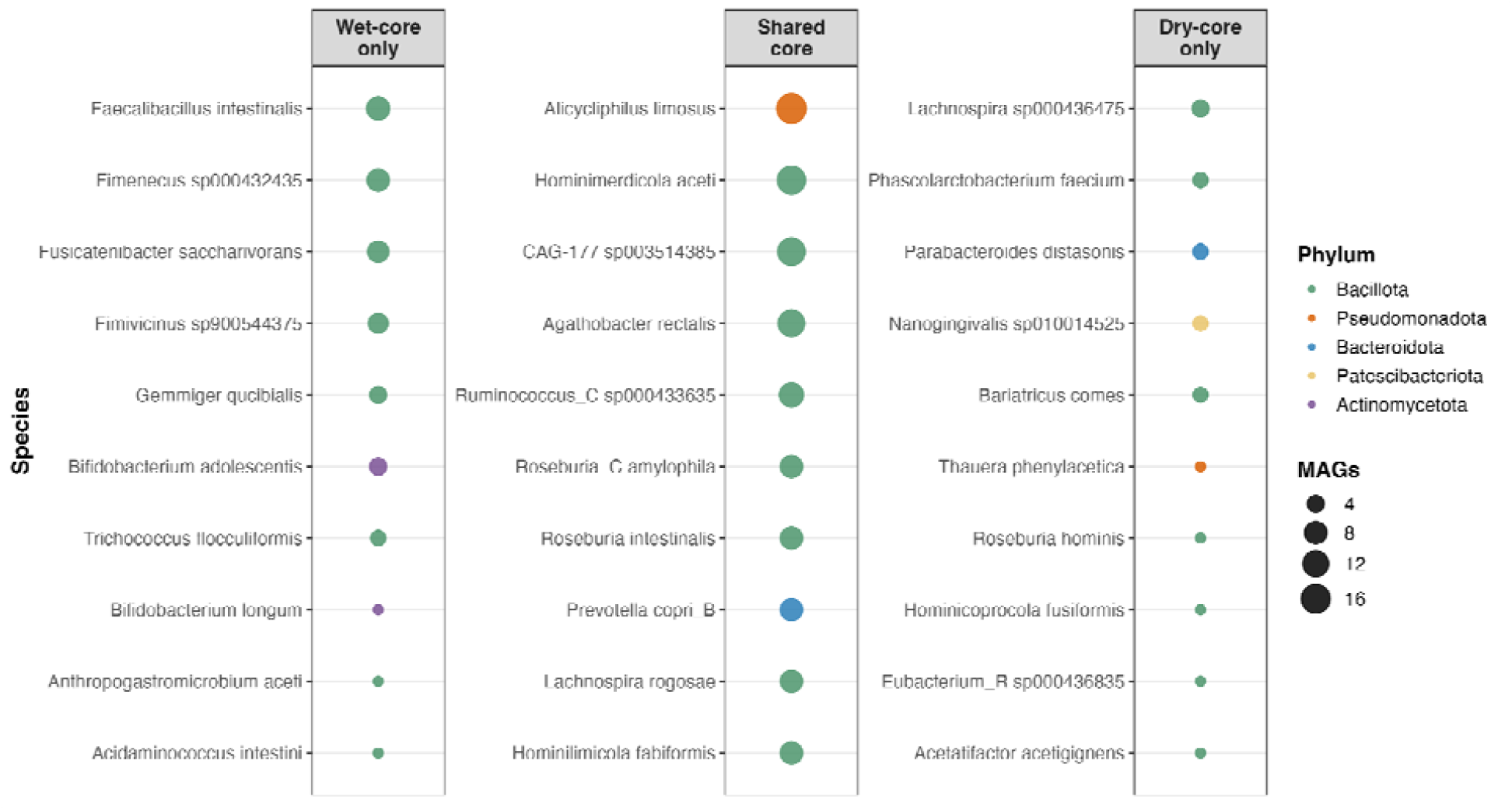
Shared and period-associated core species in MAG-derived profiles. Panels show selected wet-core-only, shared-core, and dry-core-only species. Core membership requires ≥90% prevalence within the relevant period; shared-core species meet this threshold in both periods. Bubble size indicates the number of MAGs, and color indicates phylum.

**Figure S2.**
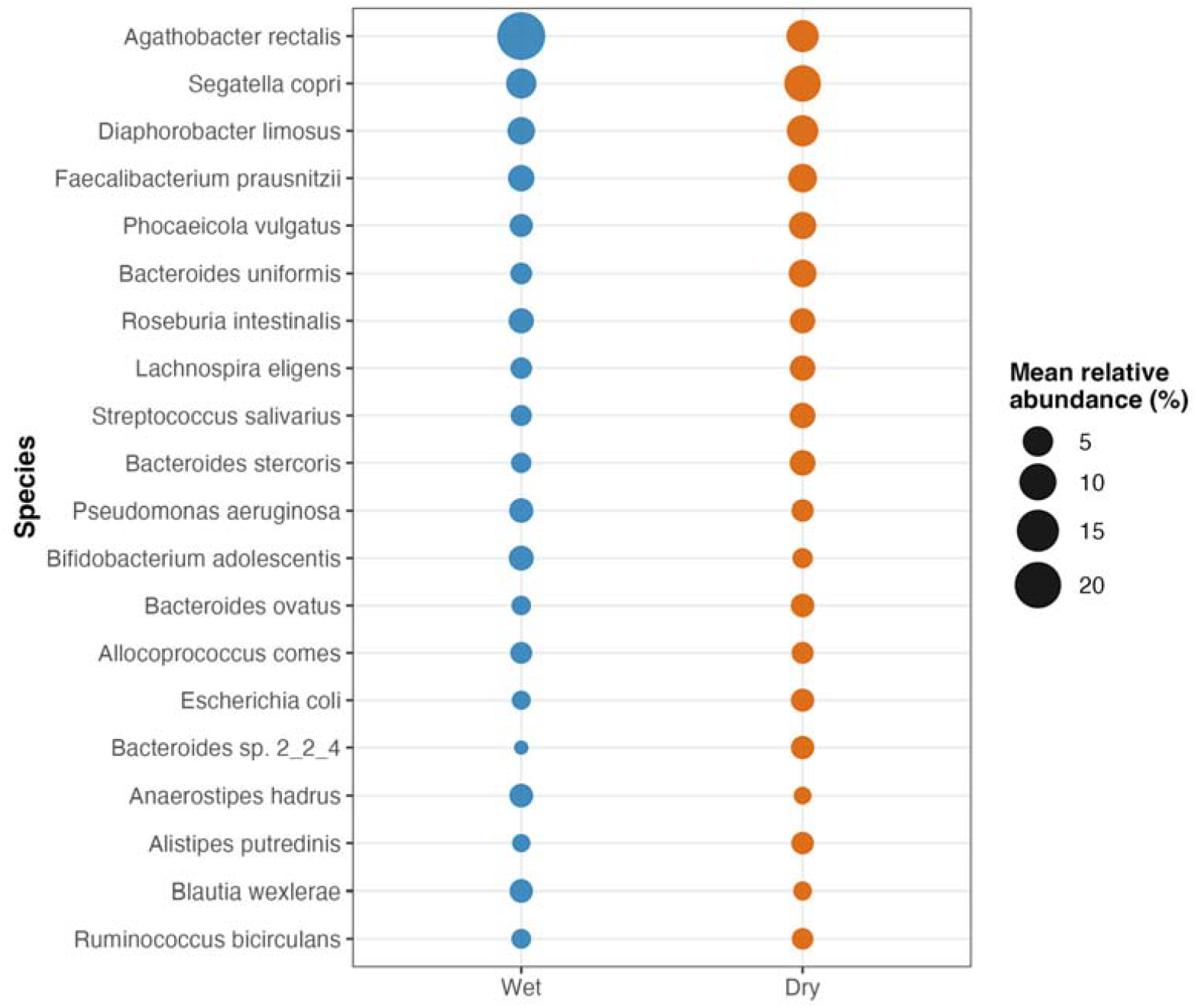
Dominant species in read-based profiles. Bubble size represents mean relative abundance (%) within the wet or dry sampling period. Each row represents a species, allowing comparison of its period-level mean relative abundance.

## References

Alcock BP, et al. 2023. CARD 2023: expanded curation, support for machine learning, and resistome prediction at the Comprehensive Antibiotic Resistance Database. Nucleic Acids Research 51:D690– D699.

Aroney STN, Newell RJP, Nissen JN, Camargo AP, Tyson GW, Woodcroft BJ. 2025. CoverM: read alignment statistics for metagenomics. Bioinformatics 41:btaf147. doi:10.1093/bioinformatics/btaf147.

Becsei Á, et al. 2024. Time-series sewage metagenomics distinguishes seasonal, human-derived and environmental microbial communities potentially allowing source-attributed surveillance. Nature Communications 15:7551. doi:10.1038/s41467-024-51957-8.

Bivins A, et al. 2022. Building-level wastewater surveillance using tampon swabs and RT-LAMP for rapid SARS-CoV-2 RNA detection. Environmental Science: Water Research & Technology 8:173–183. doi:10.1039/D1EW00496D.

Centers for Disease Control and Prevention. 2024. Kinds of E. coli. https://www.cdc.gov/ecoli/about/kinds-of-ecoli.html.

Centers for Disease Control and Prevention. 2025. About Pseudomonas aeruginosa. https://www.cdc.gov/pseudomonas-aeruginosa/about/index.html.

Centers for Disease Control and Prevention. 2026. About C. diff. https://www.cdc.gov/c-diff/about/index.html.

Chaumeil PA, Mussig AJ, Hugenholtz P, Parks DH. 2022. GTDB-Tk v2: memory friendly classification with the genome taxonomy database. Bioinformatics 38:5315–5316. doi:10.1093/bioinformatics/btac672.

Elsaghir H, Reddivari AKR. 2023. Bacteroides fragilis. StatPearls. StatPearls Publishing. https://www.ncbi.nlm.nih.gov/books/NBK553032/.

Fierer N, et al. 2022. A Metagenomic Investigation of Spatial and Temporal Changes in Sewage Microbiomes across a University Campus. mSystems 7:e00651–22. doi:10.1128/msystems.00651-22.

Gweon HS, et al. 2019. The impact of sequencing depth on the inferred taxonomic composition and AMR gene content of metagenomic samples. Environmental Microbiome 14:7. doi:10.1186/s40793-019-0347-1.

Kamali A, Ferguson D, Dowless H, Ortiz N, Mukhopadhyay R, Schember C, Lunsford R, Hutchinson J, Scherer M, Crandall J, Bauer H, Yu A, Kimura A. 2024. Outbreak of invasive Serratia marcescens among persons incarcerated in a state prison, California, USA, March 2020–December 2022. Emerging Infectious Diseases 30:S41–S48. doi:10.3201/eid3013.230801.

Langmead B, Salzberg SL. 2012. Fast gapped-read alignment with Bowtie 2. Nature Methods 9:357–359. doi:10.1038/nmeth.1923.

Li D, et al. 2015. MEGAHIT: an ultra-fast single-node solution for large and complex metagenomics assembly via succinct de Bruijn graph. Bioinformatics 31:1674–1676.

Lu J, Breitwieser FP, Thielen P, Salzberg SL. 2017. Bracken: estimating species abundance in metagenomics data. PeerJ Computer Science 3:e104. doi:10.7717/peerj-cs.104.

National Center for Biotechnology Information. n.d.-a. NCBI Pathogen Detection. National Library of Medicine. https://www.ncbi.nlm.nih.gov/pathogens/.

National Center for Biotechnology Information. n.d.-b. Citrobacter freundii. Medical Subject Headings (MeSH), National Library of Medicine. https://www.ncbi.nlm.nih.gov/mesh/68016971.

Parks DH, et al. 2015. CheckM: assessing the quality of microbial genomes recovered from isolates, single cells, and metagenomes. Genome Research 25:1043–1055.

Pruden A, Vikesland PJ, Davis BC, de Roda Husman AM. 2021. Seizing the moment: Now is the time for integrated global surveillance of antimicrobial resistance in wastewater environments. Current Opinion in Microbiology 64:91–99. doi:10.1016/j.mib.2021.09.013.

U.S. Food and Drug Administration. 2012. Bad Bug Book: Foodborne Pathogenic Microorganisms and Natural Toxins Handbook. 2nd ed. https://www.fda.gov/food/foodborne-pathogens/bad-bug-book-second-edition.

West NW, et al. 2023. Passive swab versus grab sampling for detection of SARS-CoV-2 markers in wastewater. Science of the Total Environment 889:164180. doi:10.1016/j.scitotenv.2023.164180.

Wood DE, Lu J, Langmead B. 2019. Improved metagenomic analysis with Kraken 2. Genome Biology 20:257.

World Health Organization. 2024. WHO bacterial priority pathogens list, 2024:Bacterial pathogens of public health importance to guide research, development and strategies to prevent and control antimicrobial resistance. World Health Organization. https://www.who.int/publications/i/item/9789240093461.

Wu Y-W, Simmons BA, Singer SW. 2016. MaxBin 2.0: an automated binning algorithm to recover genomes from multiple metagenomic datasets. Bioinformatics 32:605–607.

